## Supplemental data for "OTTERS: A powerful TWAS framework leveraging summary-level reference data"

#### 1. Supplemental Figures

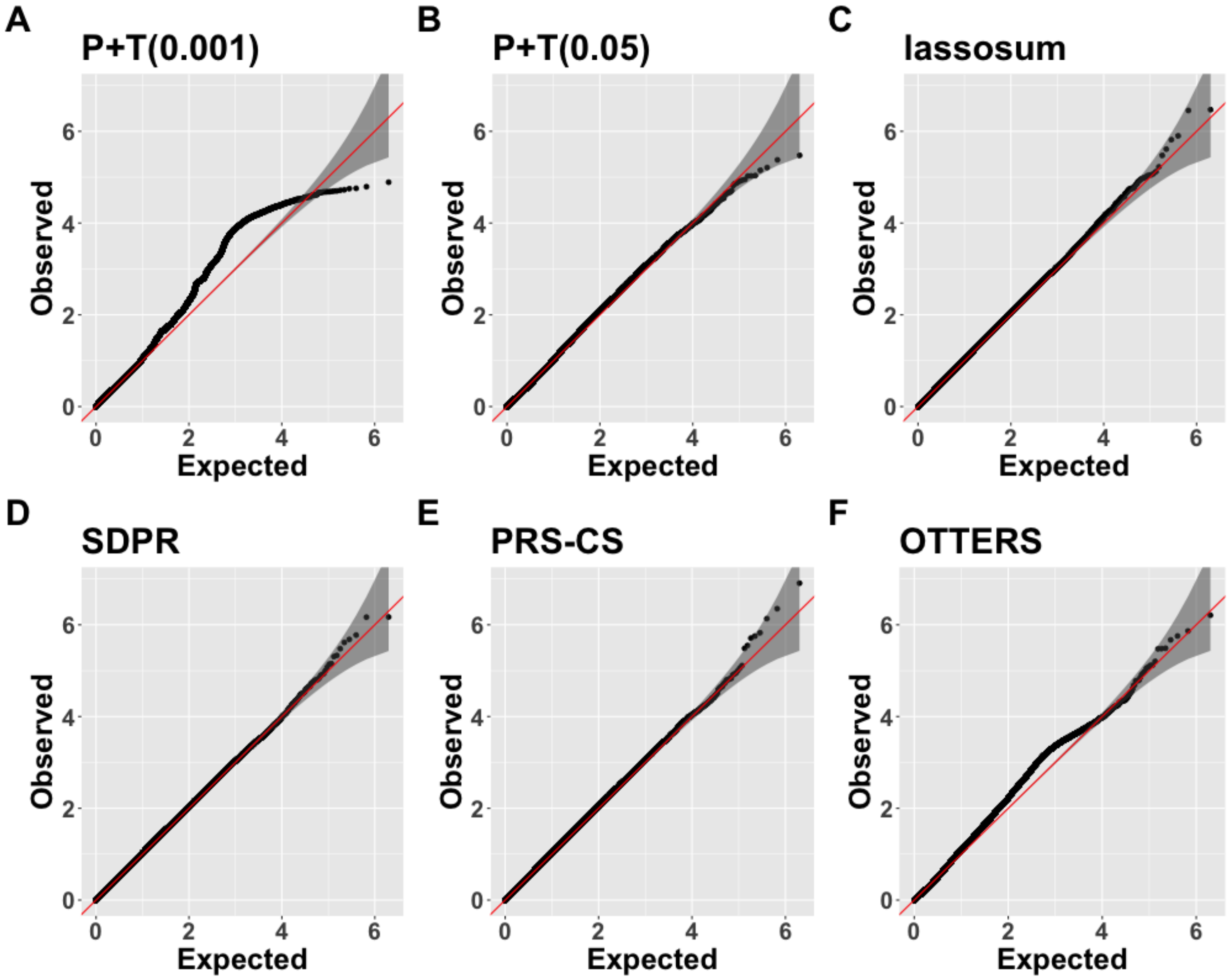


**Figure S1.** **Quantile-Quantile (QQ) plots for *P+T(0.001), P+T(0.05), lassosum, SDPR, PRS-CS*, and OTTERS under null simulations.**
Phenotype data were simulated from $N(0, 1)$. Gene expression were generated with $h_e^2 = 0.1$ and $p_{causal} = 0.001$. eQTL weights were permutated $2 \times 10^3$ times for 500 genes to perform $10^6$ null simulations of TWAS.

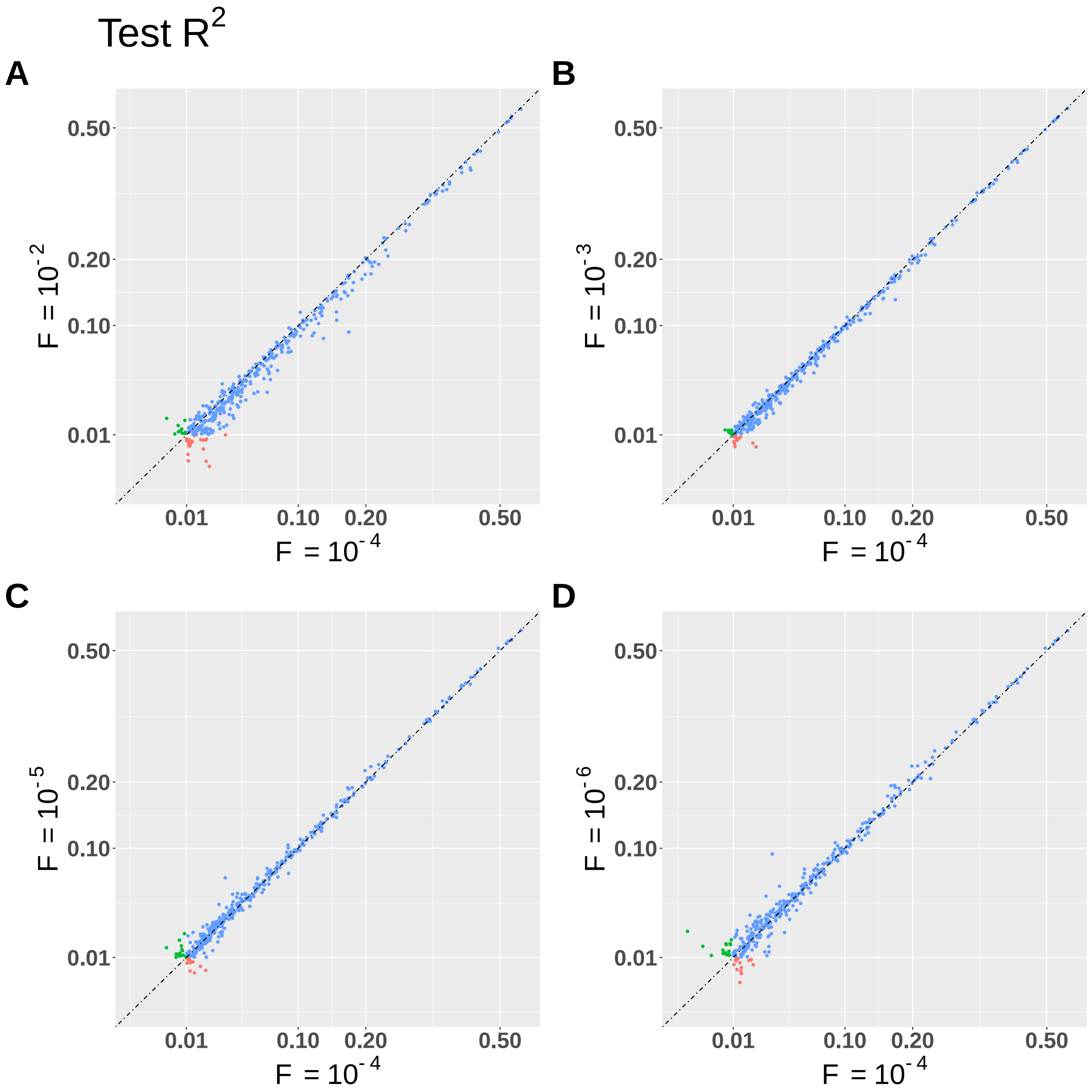


**Figure S2. Test $R^2$ in GTEx V8 samples by *PRS-CS* with $\phi$ = $(10^{-2}, 10^{-3}, 10^{-5}, 10^{-6})$ vs. $\phi = 10^{-4}$ on 590 genes on chromosome 4.**
Different colors denote whether the gene with test $R^2$> 0.01 was obtained by using both $\phi$ values (blue), or only by using one of the compared $\phi$ values (red for the $\phi$ on the x axis and green for the $\phi$ on the y axis). Genes with training $R^2 \leq 0.01$ by both methods were excluded from the plot.

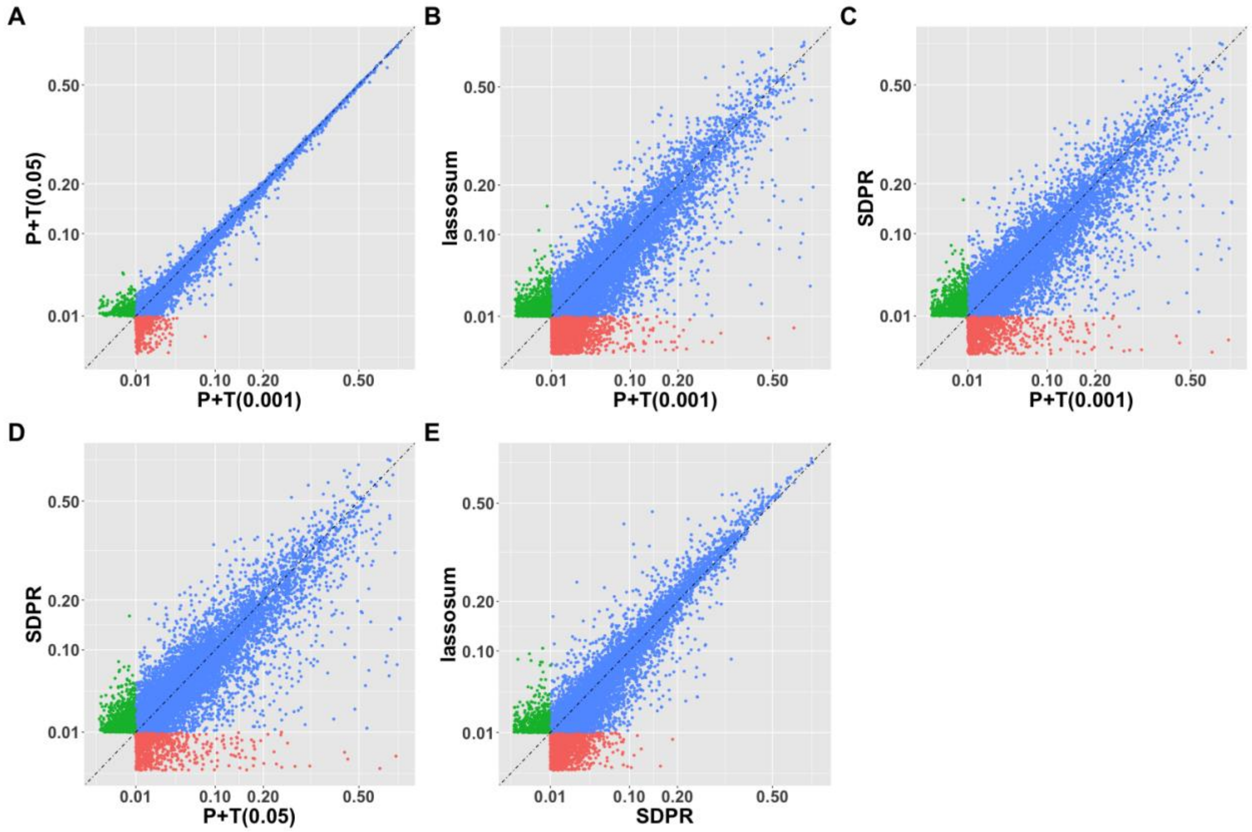


**Figure S3. Test $R^2$ in GTEx V8 whole blood samples by *P+T(0.001), P+T(0.05), lassosum,* and *SDPR.***

Different colors denoting whether the gene with test $R^2 > 0.01$ was obtained by both methods (blue), or only by one of the pair-wise compared methods (red for the method on x axis and green for the method on y axis).

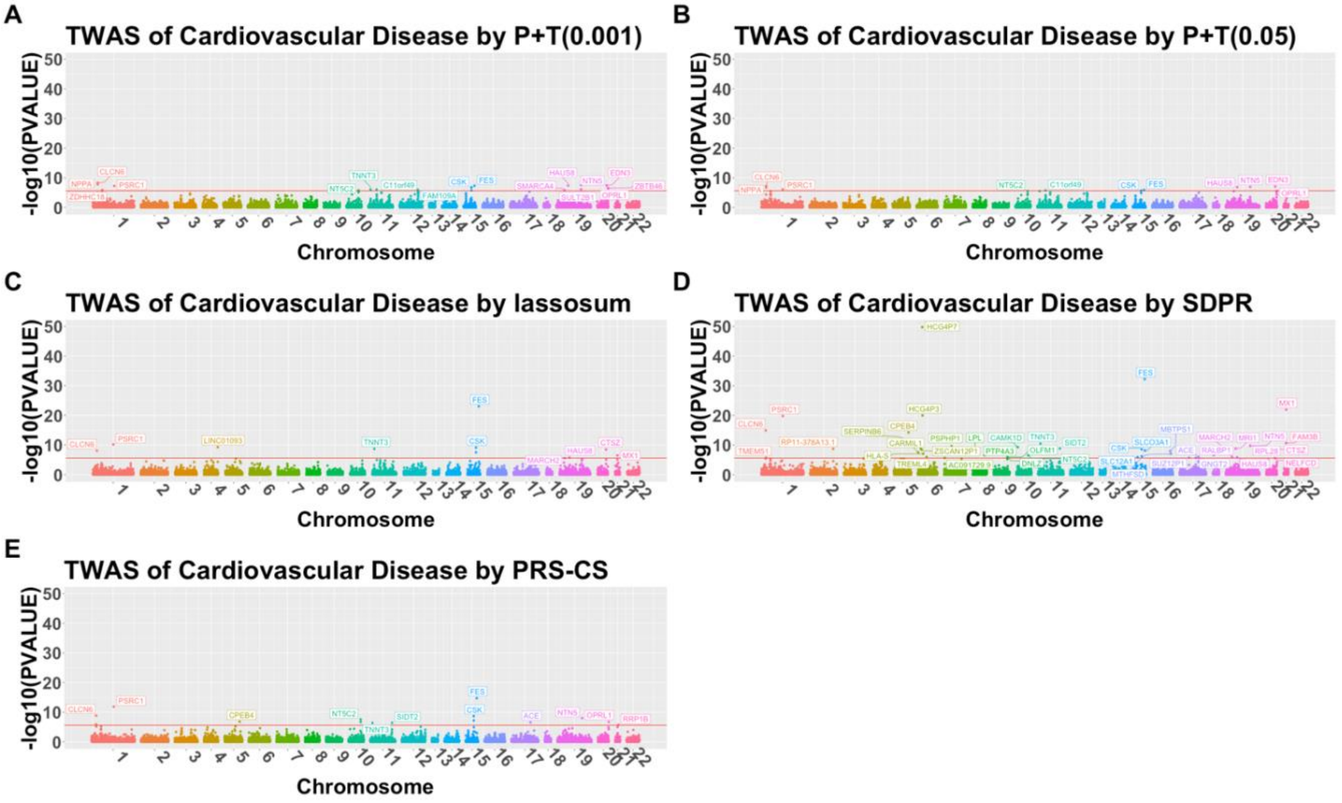


**Figure S4. Manhattan plots of TWAS of cardiovascular disease by *P+T(0.001)* (A), *P+T(0.05)* (B), *lassosum* (C), *SDPR* (D), and *PRS-CS* (E) with independently significant TWAS risk genes being labelled.**

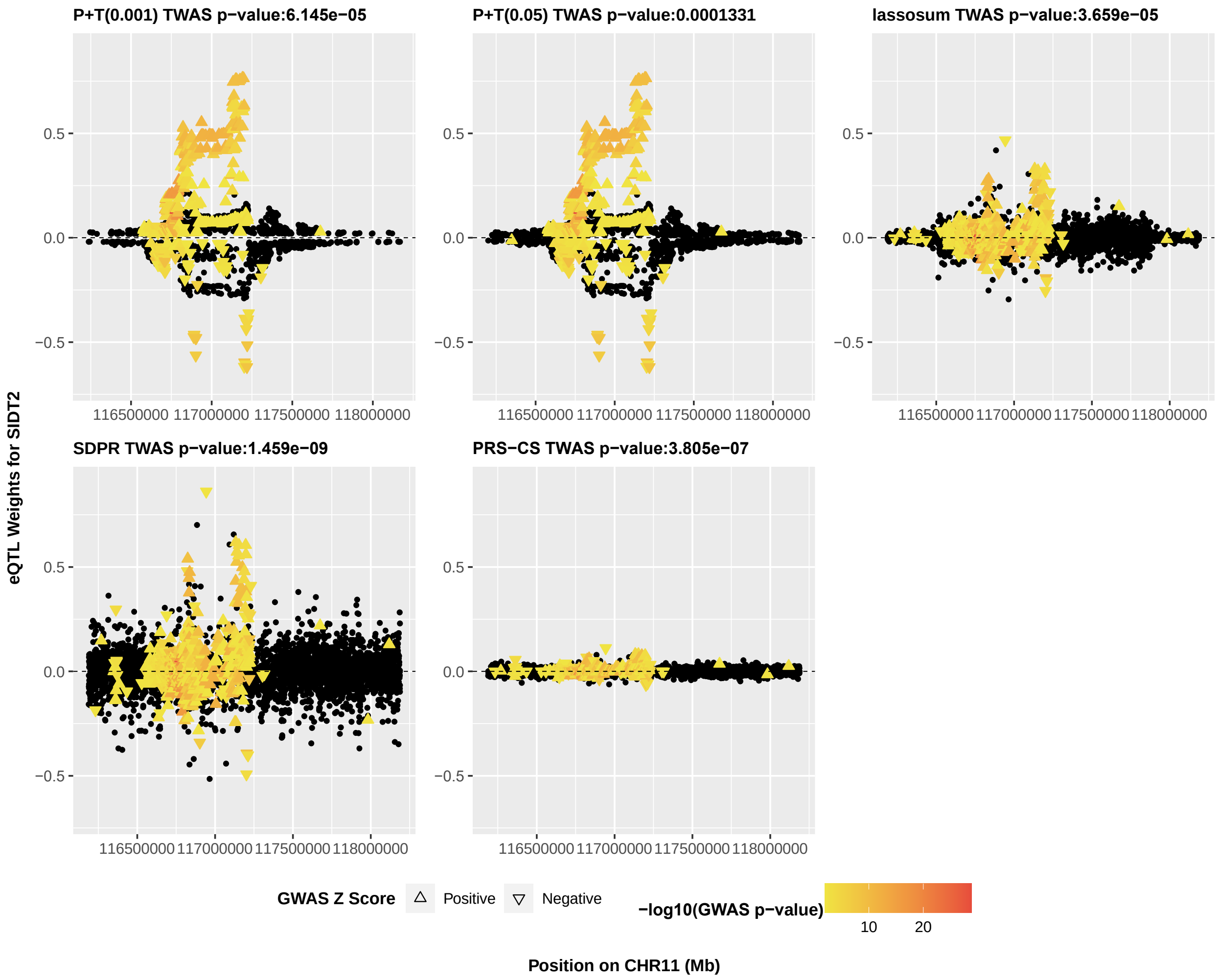


**Figure S5. Cis-eQTL weights estimated by individual methods for gene *SIDT2*, with color coded with respect to $-log_{10}$(p-value) from the UKBB GWAS summary statistics and shape coded with respect to the direction of GWAS Z-score statistics.**

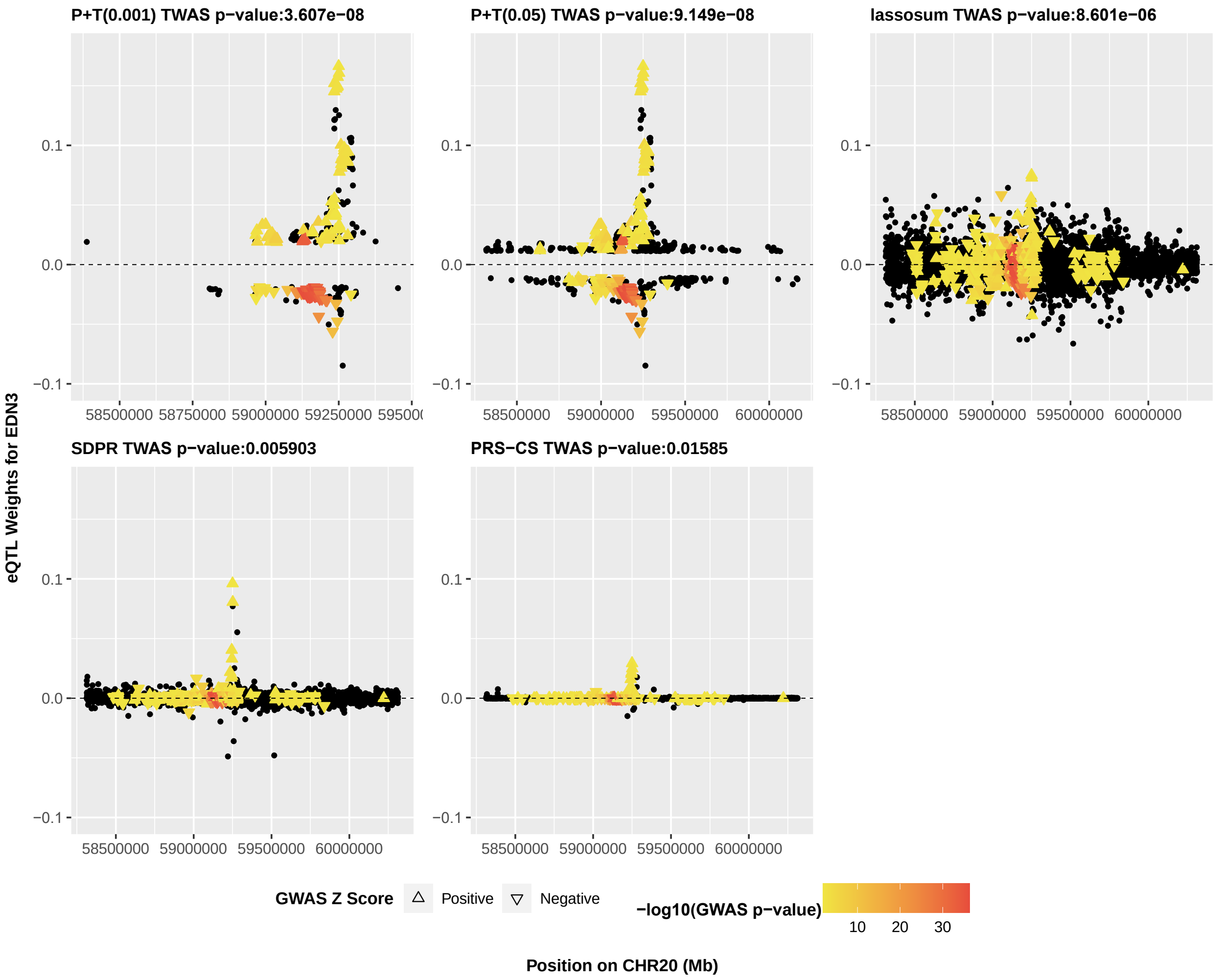


**Figure S6. Cis-eQTL weights estimated by individual methods for gene *EDN3*, with color coded with respect to $-log_{10}$(p-value) from the UKBB GWAS summary statistics and shape coded with respect to the direction of GWAS Z-score statistics.**

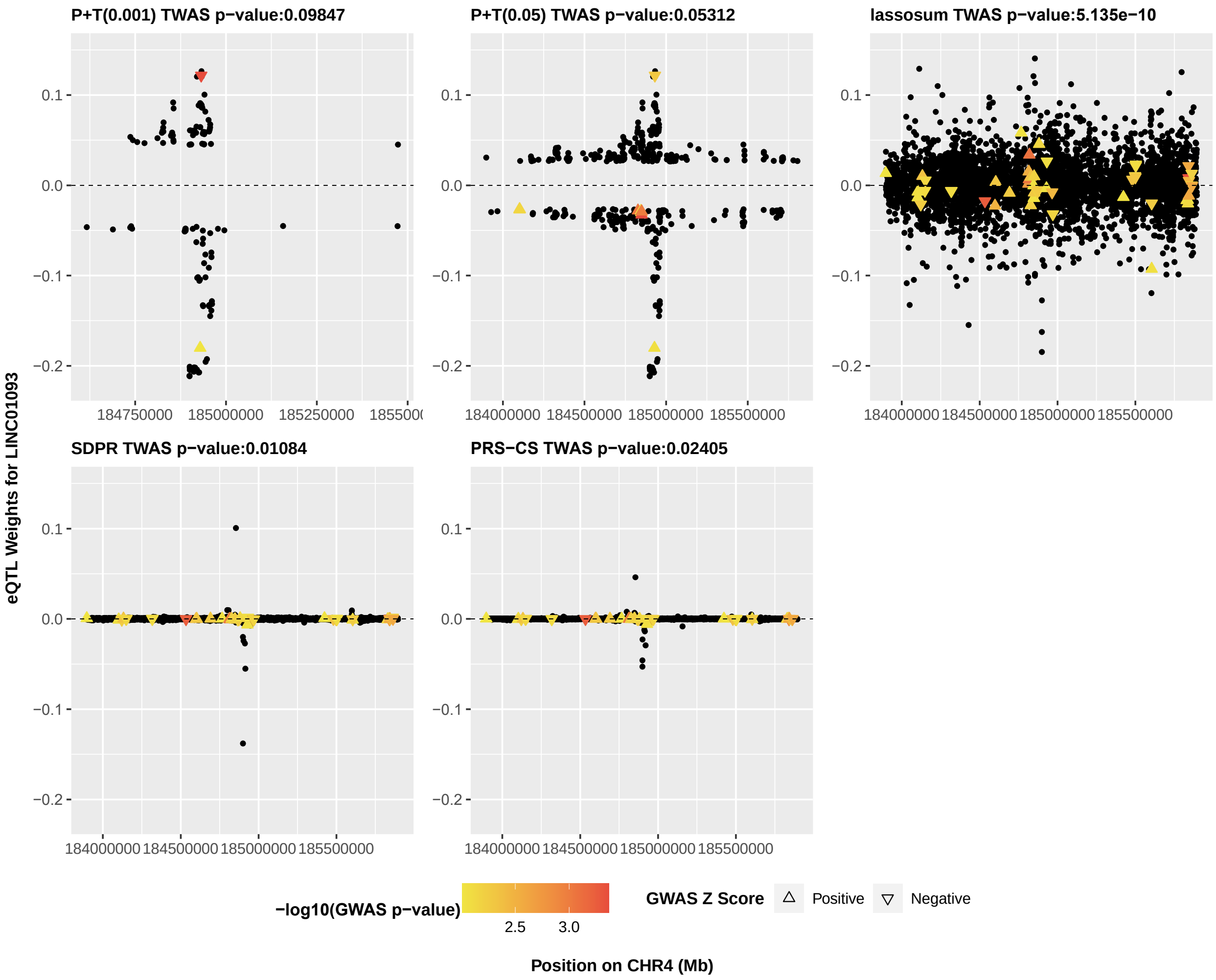


**Figure S7. cis-eQTL weights estimated by individual methods for gene LINC01093, with color coded with respect to $-log_{10}$(p-value) from the UKBB GWAS summary statistics and shape coded with respect to the direction of GWAS Z-score statistics.**

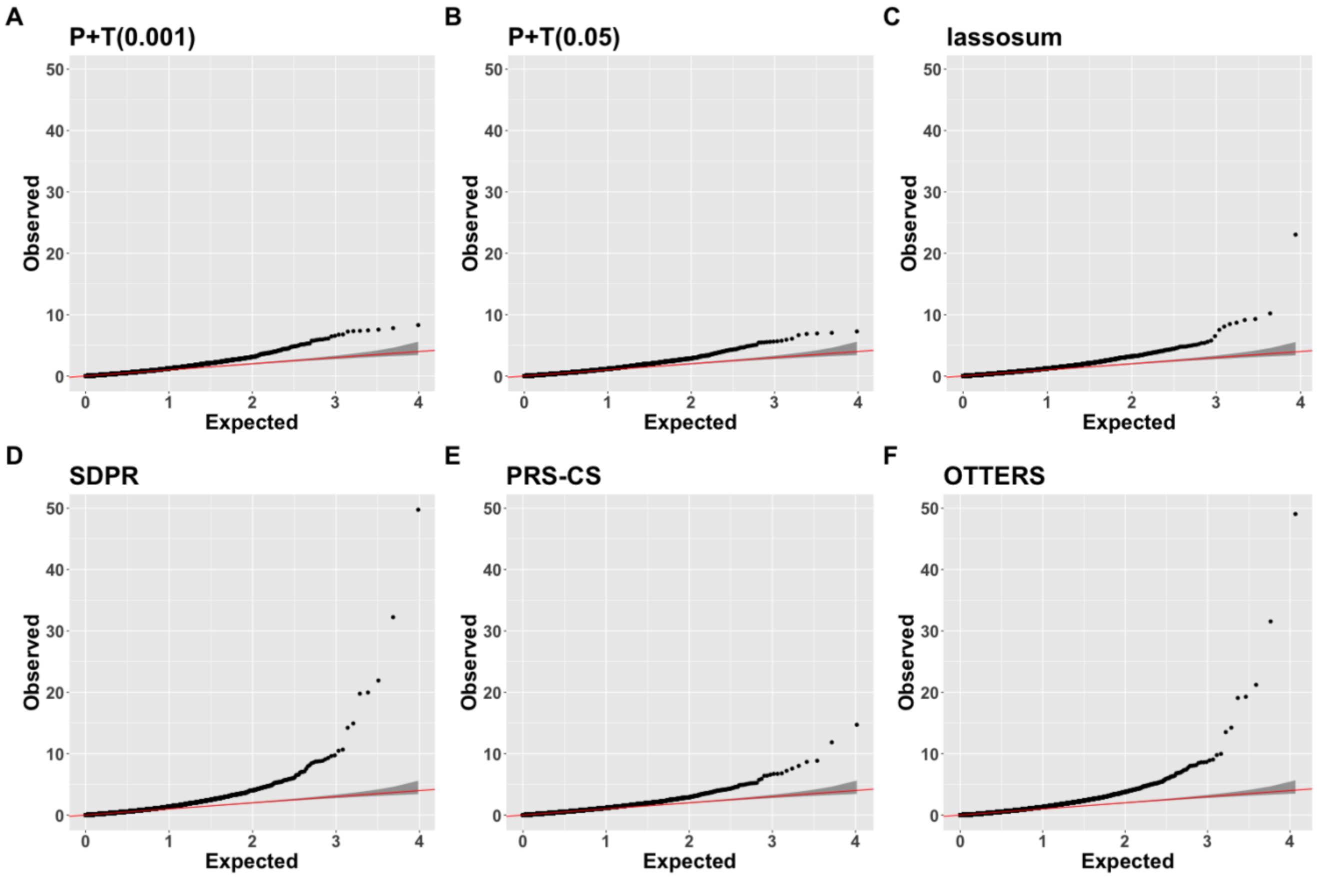


**Figure S8. Quantile-Quantile plots of TWAS p-values of cardiovascular disease by *P+T(0.001)* (A), *P+T(0.05)* (B), *lassosum* (C), *SDPR* (D), *PRS-CS* (E), and OTTERS**

**A.**

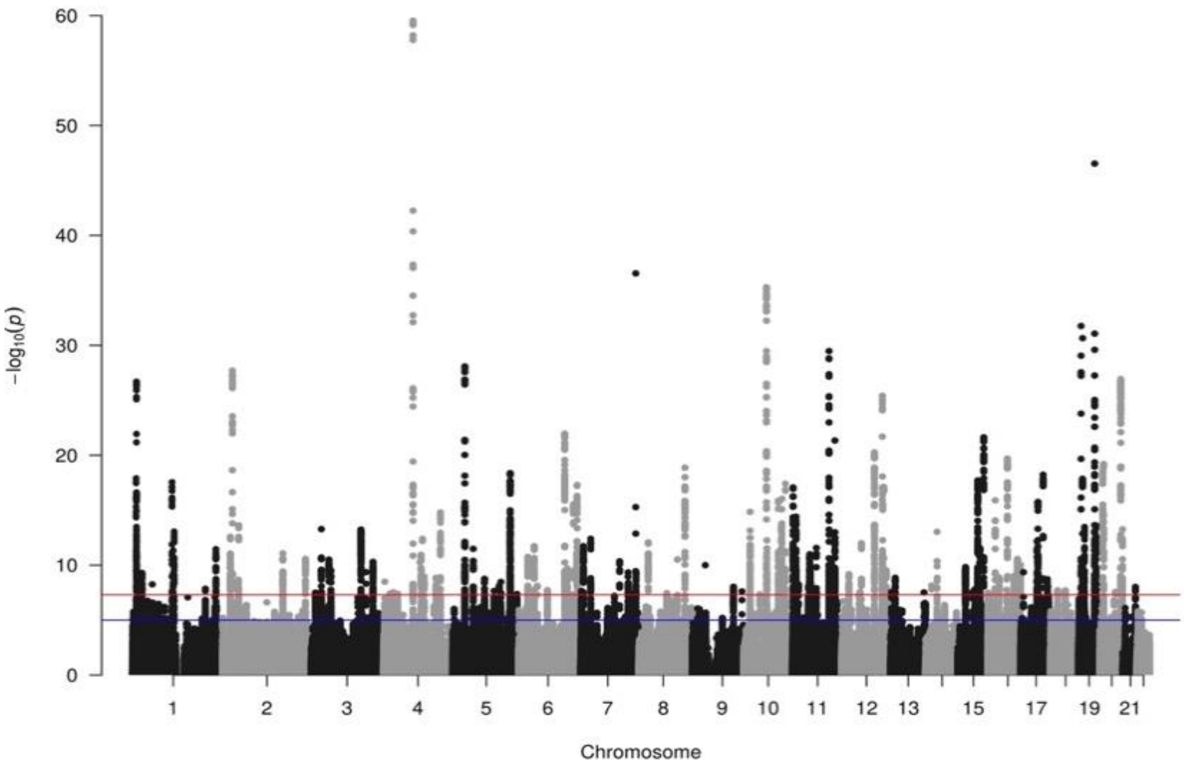


**B.**

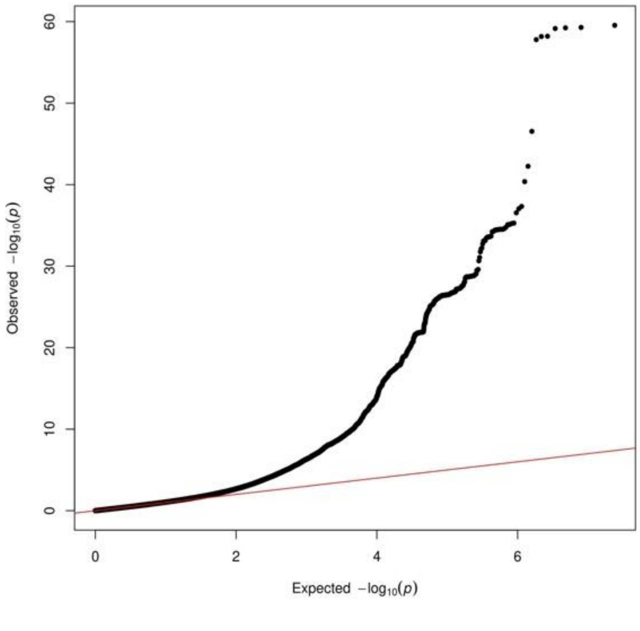


**Figure S9. Manhattan Plot (A) and Quantile-Quantile plot (B) of genomic control corrected UK Biobank GWAS summary results of cardiovascular disease.**

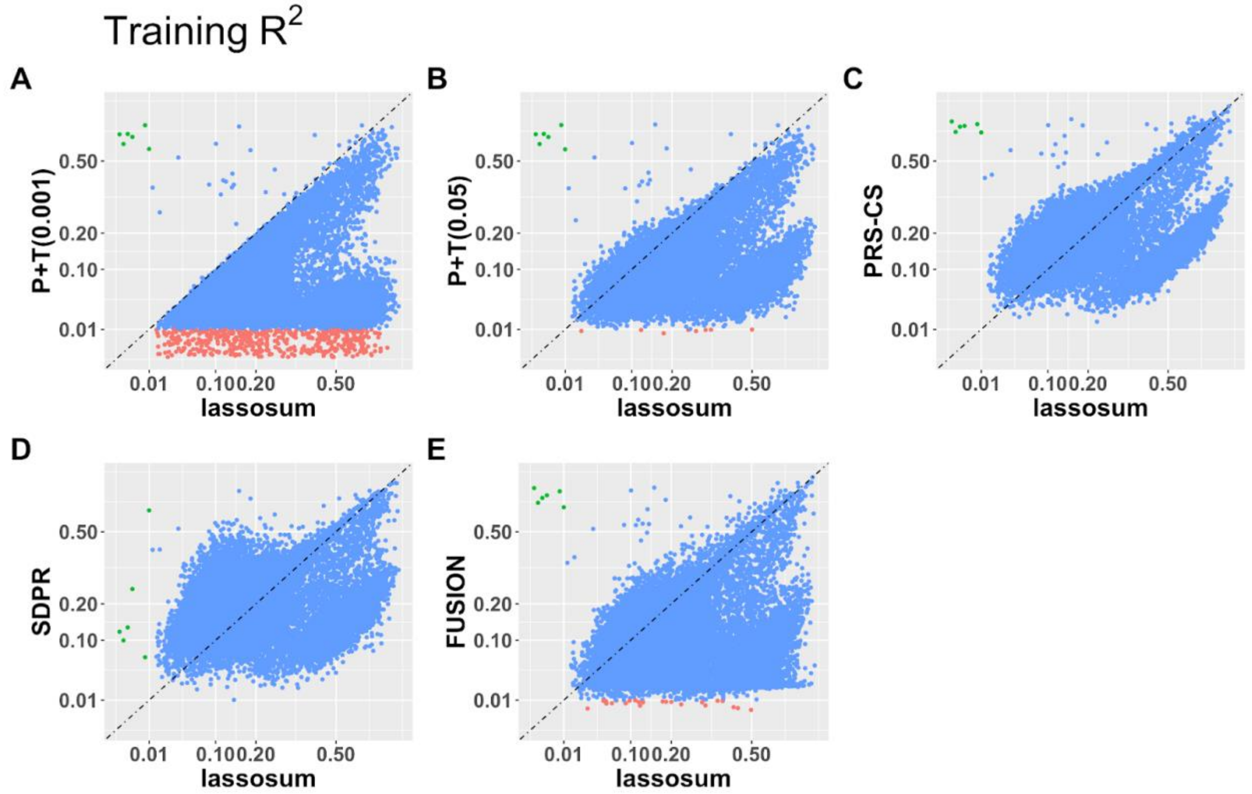


**Figure S10. Training $R^2$ in GTEx V8 whole blood samples by *lassosum versus P+T(0.001), P+T(0.05), PRS-CS*, *SDPR*, and FUSION.**

Training $R^2$ by *lassosum* versus *P+T(0.001)* (A), *P+T(0.05)* (B), *PRS-CS* (C), *SDPR* (D), and *FUSION* (E) with 574 GTEx V8 training samples, with different colors denoting whether the imputation $R^2 > 0.01$ only by *lassosum* (red), only by the y axis method (green), or both methods (blue). Genes with $R^2 \leq 0.01$ by both methods were excluded from the plot.

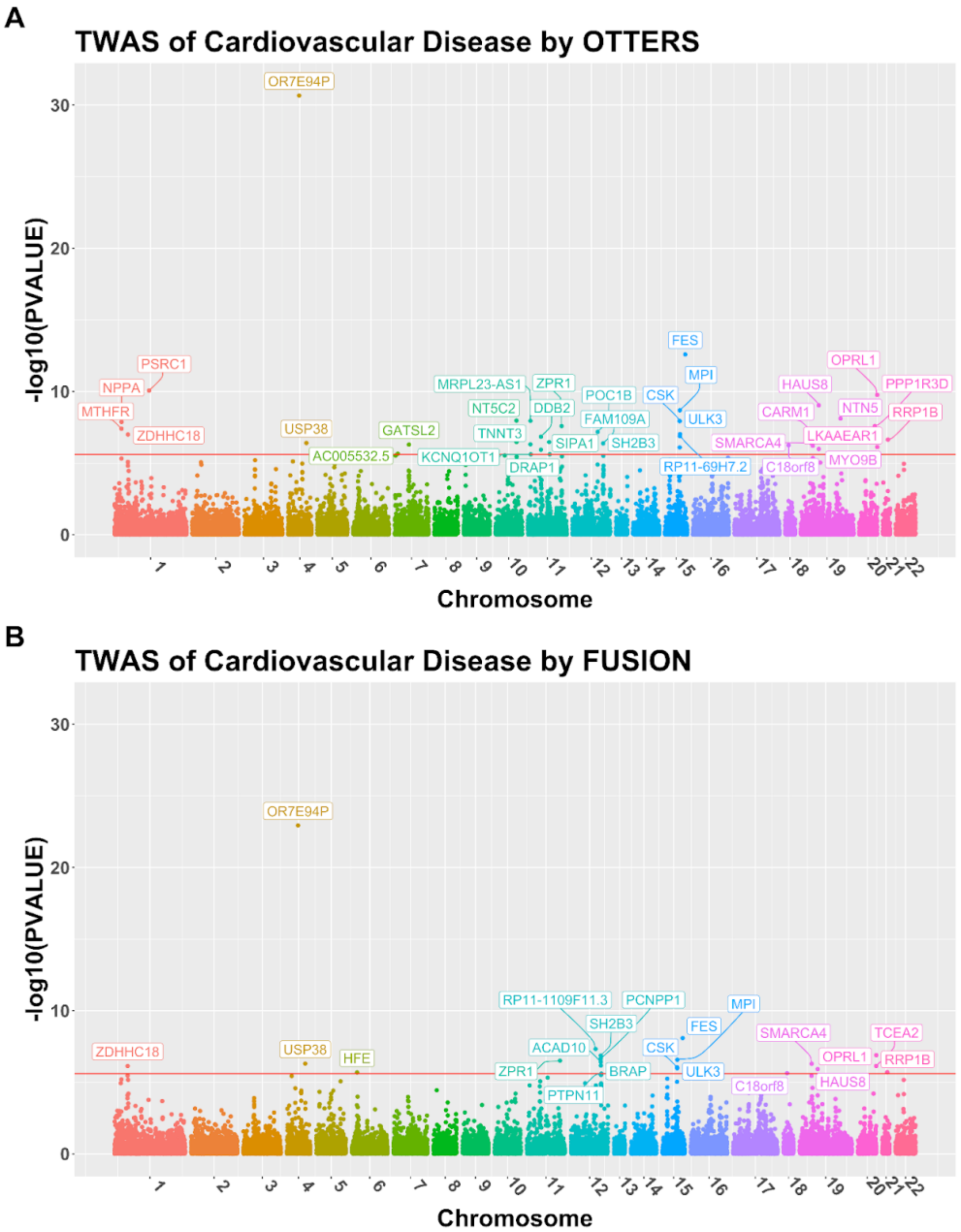


**Figure S11. Manhattan plot of TWAS results by OTTERS (A) and FUSION (B) using eQTL data from GTEx V8 whole blood samples. Independently significant TWAS risk genes are labeled.**

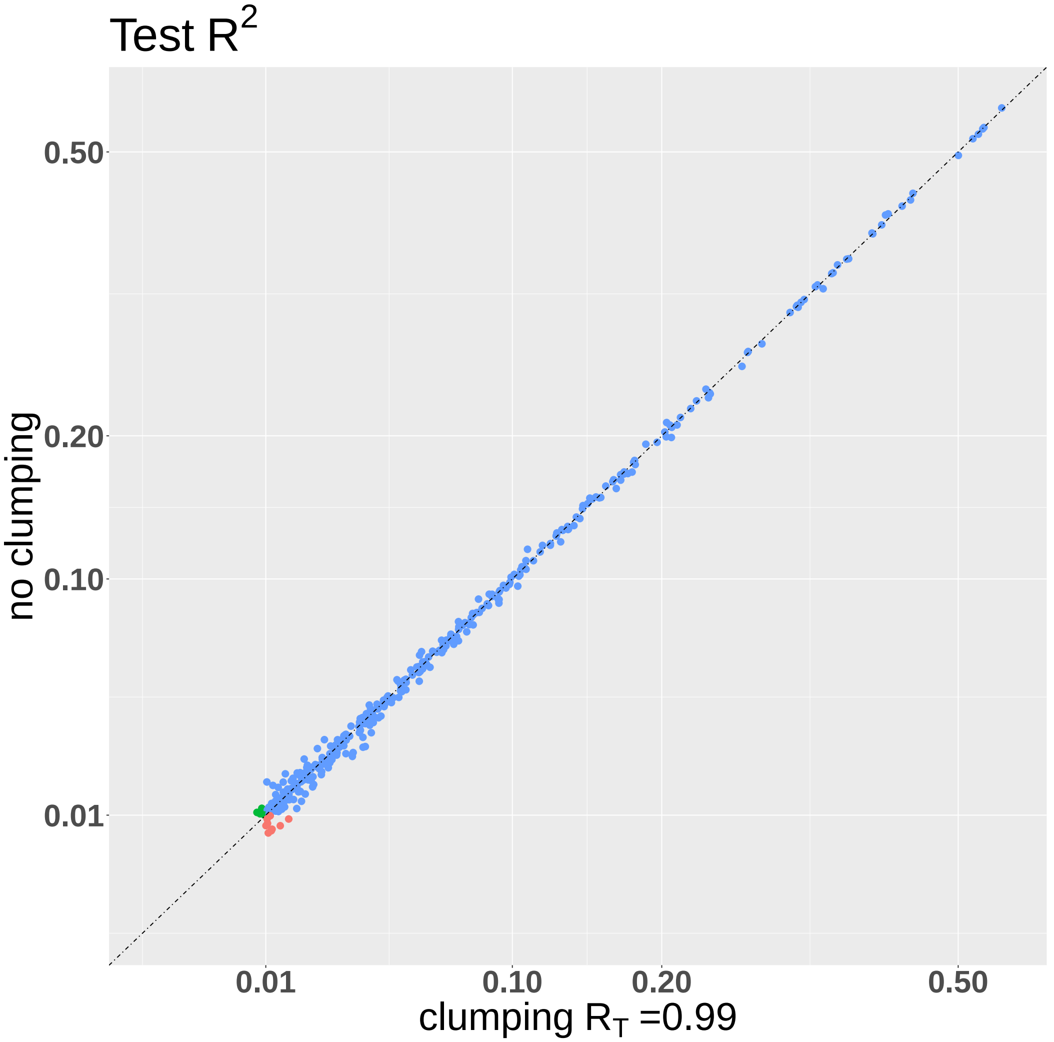


**Figure S12. Test $R^2$ in GTEx V8 samples by *PRS-CS* with LD-clumping ($R_T$ = 0.99) versus no clumping.**

Test $R^2$ in 315 GTEx V8 test samples by *PRS-CS* with LD-clumping ($R_T$ = 0.99) versus no clumping, with different colors denoting whether the imputation $R^2$ > 0.01 only by clumping (red), only by no clumping (green), or both (blue). Genes with $R^2 \leq 0.01$ by both methods were excluded from the plot.

### 2. Supplemental Tables

**Table S1. TWAS type I errors under null simulation studies with $h_e^2 = 0.1$ and $p_{causal} = 0.001$ by 5 individual methods *P+T(0.001), P+T(0.05), lassosum, SDPR, PRS-CS*, and OTTERS.**

| Significance Level | P+T(0.001) | P+T(0.05) | lassosum | SDPR | PRS-CS | OTTERS |
|---|---|---|---|---|---|---|
| $\mathbf{1.0 \times 10^{-2}}$ | $1.71 \times 10^{-2}$ | $1.25 \times 10^{-2}$ | $1.12 \times 10^{-2}$ | $1.08 \times 10^{-2}$ | $1.12 \times 10^{-2}$ | $1.60 \times 10^{-2}$ |
| $\mathbf{1.0 \times 10^{-4}}$ | $7.13 \times 10^{-5}$ | $9.60 \times 10^{-5}$ | $1.32 \times 10^{-4}$ | $9.70 \times 10^{-5}$ | $1.13 \times 10^{-4}$ | $9.30 \times 10^{-5}$ |
| $\mathbf{2.5 \times 10^{-6}}$ | 0 | 0 | $5.00 \times 10^{-6}$ | $5.00 \times 10^{-6}$ | $5.00 \times 10^{-6}$ | $4.00 \times 10^{-6}$ |

**Table S2. Independent significant TWAS genes of cardiovascular disease identified by individual training methods (*P+T(0.001), P+T(0.05), lassosum, SDPR, PRS-CS*) but not by OTTERS.**

| | CHR | GENE | P-VALUE | OTTERS P-VALUE |
|---|---|---|---|---|
| **P+T (0.001)** | 1 | ZDHHC18 | 1.2602E-06 | 3.8725E-06 |
| | 12 | FAM109A | 1.1125E-06 | 3.8897E-06 |
| | 19 | SMARCA4 | 1.4672E-06 | 5.0821E-06 |
| **lassosum** | 19 | MARCH2 | 2.6164E-06 | 4.0247E-06 |
| **SDPR** | 1 | TMEM51 | 1.8037E-06 | 9.0175E-06 |
| | 6 | TREML4 | 9.6488E-07 | 4.8197E-06 |
| | 7 | AC091729.9 | 2.3835E-06 | 1.191E-05 |
| | 9 | OLFM1 | 1.3473E-06 | 6.736E-06 |
| | 9 | DNLZ | 1.8569E-06 | 9.2841E-06 |
| | 15 | SLC12A1 | 1.2396E-06 | 6.1852E-06 |
| | 17 | SUZ12P1 | 1.9124E-06 | 9.4828E-06 |
| | 17 | GNGT2 | 2.0687E-06 | 9.8317E-06 |
| | 19 | MARCH2 | 1.1691E-06 | 4.0247E-06 |
| | 20 | NELFCD | 2.8779E-06 | 1.4186E-05 |
| **PRS-CS** | 21 | RRP1B | 1.8223E-06 | 6.9344E-06 |

**Table S3. Independent TWAS risk genes of cardiovascular disease identified by OTTERS using eQTL summary statistics from GTEx V8 whole blood samples.**

| **CHR** | **ID** | OTTERS | **FUSION** | **P+T(0.001)** | **P+T(0.05)** | **lassosum** | **SDPR** | **PRScs** |
|---|---|---|---|---|---|---|---|---|
| 1 | MTHFR | **3.99E-08** | 1.12E-03 | **8.05E-09** | **1.11E-06** | 7.17E-05 | 1.06E-03 | 7.95E-05 |
| 1 | NPPA | **1.33E-08** | 6.35E-06 | **1.19E-08** | **3.52E-09** | **1.39E-07** | 5.71E-04 | 5.94E-04 |
| 1 | ZDHHC18 | **1.02E-07** | **7.35E-07** | **1.82E-07** | **5.27E-07** | **2.67E-08** | **9.34E-07** | **3.18E-07** |
| 1 | PSRC1 | **8.75E-11** | 1.70E-04 | **1.91E-11** | **1.61E-06** | **1.23E-09** | **3.11E-10** | **1.45E-09** |
| 4 | OR7E94P | **2.14E-31** | **1.16E-23** | **4.28E-32** | 2.53E-04 | **2.16E-10** | 2.80E-05 | **1.35E-11** |
| 4 | USP38 | **3.98E-07** | **4.99E-07** | 2.85E-05 | 1.09E-04 | **8.12E-08** | 7.85E-06 | 1.41E-05 |
| 7 | AC005532.5 | **2.26E-06** | 1.03E-03 | **4.56E-07** | 4.88E-05 | 3.89E-04 | 1.17E-03 | 7.48E-04 |
| 7 | GATSL2 | **5.12E-07** | 1.69E-02 | **7.59E-07** | **1.18E-07** | 1.70E-03 | 3.45E-03 | 7.02E-04 |
| 10 | NT5C2 | **1.09E-08** | 2.42E-04 | **2.69E-09** | **1.76E-06** | **1.17E-08** | 1.03E-03 | 6.62E-05 |
| 11 | TNNT3 | **4.89E-07** | 1.65E-05 | **1.01E-07** | 1.16E-05 | 7.80E-06 | 2.72E-05 | 3.02E-05 |
| 11 | MRPL23-AS1 | **1.13E-08** | 2.48E-01 | **2.26E-09** | 1.21E-05 | 8.89E-05 | 7.11E-02 | 8.61E-04 |
| 11 | KCNQ1OT1 | **2.47E-06** | 7.37E-03 | **5.09E-07** | 1.32E-04 | 1.80E-05 | 1.85E-02 | 4.39E-03 |
| 11 | DDB2 | **1.46E-07** | 2.14E-04 | **1.00E-07** | 3.52E-06 | **4.17E-08** | 4.10E-04 | 1.18E-03 |
| 11 | SIPA1 | **3.42E-07** | 4.77E-06 | **1.24E-07** | 3.49E-05 | **1.72E-07** | 4.31E-05 | **1.45E-06** |
| 11 | DRAP1 | **2.46E-06** | 9.41E-02 | **4.92E-07** | 4.16E-02 | 1.88E-01 | 4.20E-02 | 9.94E-02 |
| 11 | ZPR1 | **2.64E-08** | **3.11E-07** | **9.47E-07** | **5.34E-09** | 2.06E-05 | 3.44E-06 | **1.09E-06** |
| 12 | POC1B | **6.76E-08** | 8.87E-05 | **1.35E-08** | 5.55E-05 | 1.19E-05 | 9.13E-04 | 1.08E-04 |
| 12 | FAM109A | **4.25E-07** | 1.44E-05 | **3.58E-07** | **3.05E-07** | 1.04E-03 | **4.69E-07** | **2.82E-07** |
| 12 | SH2B3 | **4.27E-07** | **1.37E-07** | **2.01E-07** | **5.86E-07** | 4.36E-04 | **9.32E-07** | **2.52E-07** |
| 15 | CSK | **1.18E-08** | **9.12E-07** | **3.12E-07** | **4.20E-07** | 5.53E-01 | **2.28E-08** | **2.68E-09** |
| 15 | ULK3 | **1.18E-08** | **1.05E-06** | **8.28E-09** | **1.05E-07** | 6.39E-01 | **4.27E-09** | **1.68E-08** |
| 15 | MPI | **2.08E-09** | **2.69E-07** | **3.61E-08** | **1.58E-07** | 4.47E-01 | **6.20E-10** | **1.32E-09** |
| 15 | RP11-69H7.2 | **9.54E-08** | 5.12E-03 | **1.91E-08** | 2.66E-05 | 1.39E-04 | 6.35E-05 | 2.78E-05 |
| 15 | FES | **2.61E-13** | **8.18E-09** | **8.34E-14** | **1.16E-08** | **4.48E-12** | **4.35E-12** | **1.48E-13** |
| 18 | C18orf8 | **5.71E-07** | **2.32E-06** | 9.16E-06 | **1.87E-06** | **5.94E-07** | **1.99E-07** | **7.05E-07** |
| 19 | CARM1 | **7.43E-08** | 7.53E-02 | 3.12E-06 | **1.49E-08** | 6.60E-04 | 2.52E-03 | 1.87E-03 |
| 19 | SMARCA4 | **6.24E-07** | **5.14E-07** | **4.26E-07** | **9.35E-07** | **2.17E-07** | 1.34E-01 | 1.37E-03 |
| 19 | HAUS8 | **9.29E-10** | **1.20E-06** | **1.56E-09** | **1.85E-09** | **2.38E-10** | **5.11E-07** | **9.97E-08** |
| 19 | MYO9B | **1.04E-06** | 8.26E-05 | **2.15E-07** | 9.87E-06 | 4.28E-04 | 3.78E-03 | 2.97E-05 |
| 19 | NTN5 | **7.77E-09** | 9.62E-06 | **1.55E-09** | 3.95E-06 | 5.29E-06 | 4.78E-03 | 2.56E-04 |
| 20 | PPP1R3D | **2.63E-08** | 1.09E-01 | 3.21E-05 | **5.25E-09** | 3.51E-01 | 9.16E-04 | 8.22E-03 |
| 20 | OPRL1 | **1.74E-10** | **1.28E-07** | **4.16E-08** | **8.25E-07** | **3.53E-11** | **5.25E-09** | **4.55E-09** |
| 20 | LKAAEAR1 | **7.50E-07** | 3.00E-01 | **1.50E-07** | 8.47E-04 | 2.68E-01 | 1.14E-02 | 1.04E-03 |
| 21 | RRP1B | **2.35E-07** | **2.00E-06** | **5.73E-08** | **4.56E-07** | **6.27E-07** | 8.62E-03 | 1.32E-04 |

**Table S4. Computational time (minutes) and memory usage (Gigabytes in the parenthesis) of different training methods for genes with different number of test eQTLs after LD-clumping with $R_T$ = 0.99.**

For each group with the number of test eQTLs <2000, from 2000 to 3000, from 3000 to 4000, and >4000, the reported computational time and memory usage were the average time and memory per gene used for 10 randomly selected genes on Chromosome 4.

| **# Test eQTLs**[a] | P+T | lassosum | SDPR | PRS-CS |
|---|---|---|---|---|
| **<2000** | 0.003 (0.0015) | 0.16 (0.025) | 3.56 (0.024) | 4.18 (0.525) |
| **2000 - 3000** | 0.003 (0.0012) | 0.17 (0.027) | 9.07 (0.065) | 7.68 (0.416) |
| **3000 - 4000** | 0.005 (0.0013) | 0.20 (0.029) | 7.96 (0.06) | 35.14 (0.573) |
| **>4000** | 0.005 (0.0008) | 0.54 (0.035) | 30.89 (0.173) | 89.945 (0.577) |

a: number of eQTLs per gene after LD-clumping ($R_T$ = 0.99).

### 3. Supplemental Methods

#### 3.1 OTTERS Stage I with Summary-level Reference Data

OTTERS will train eQTL effect sizes $\hat{\boldsymbol{w}}$ using the following GReX imputation model (i.e., a multivariate regression model assuming additive genetic effect sizes):

$$\boldsymbol{E}_g = \boldsymbol{X}\boldsymbol{w} + \boldsymbol{\epsilon},\ \ \boldsymbol{\epsilon} \sim N(0, \sigma_\epsilon^2 \boldsymbol{I}), \qquad \text{(Equation S1)}$$

based on summary-level reference data that contain marginal eQTL effect sizes ($\tilde{w}_j, j = 1, \dots, m$) and p-values from the following single variant regression model:

$$\boldsymbol{E}_g = \boldsymbol{X}_j w_j + \boldsymbol{\epsilon},\ \ \boldsymbol{\epsilon} \sim N(0, \sigma_\epsilon^2 \boldsymbol{I}),\ j\ =\ 1, \dots, m. \qquad \text{(Equation S2)}$$

Here, $\boldsymbol{E}_g$ is a vector representing gene expression levels of gene $g$, $\boldsymbol{X}_j$ is an $n \times 1$ vector of genotype data for genetic variant $j$, $\boldsymbol{X}$ is an $n \times m$ matrix of genotype data of SNP predictors proximal or within gene $g$, $\boldsymbol{w}$ is the eQTL effect-size vector, $n$ is the sample size, and $\boldsymbol{\epsilon}$ is the error term.

Assuming $\boldsymbol{E_g}$ and the column of $\boldsymbol{X}$ to be standardized, $\boldsymbol{E}_g^T\boldsymbol{E}_g = n$, we can show that the marginal least squared eQTL effect size estimates from Equation S2 is $\tilde{\boldsymbol{w}} = \boldsymbol{X}^T\boldsymbol{E}_g/n$ and that the LD correlation matrix is $\boldsymbol{R} = \boldsymbol{X}^T\boldsymbol{X}/n$. OTTERS uses four summary-data based polygenic risk score (PRS) methods in this step — P-value Thresholding with linkage disequilibrium (LD) clumping (*P*+*T*)[1], frequentist LASSO[2] regression based method *lassosum*[3], Bayesian Dirichlet Process Regression model[4] based method *SDPR*[5], and Bayesian multivariate regression model based method *PRS-CS*[6]. The *P*+*T* method is described in the main text. Here, we describe details of the other three methods.

##### 3.1.1 Frequentist *lassosum*

The LASSO[2] estimates $\hat{\boldsymbol{w}}$ by minimizing the following loss function with a penalty parameter $\lambda$,

$$f(\boldsymbol{w}) = \frac{\boldsymbol{E}_g^T \boldsymbol{E}_g}{n} + \boldsymbol{w}^T \boldsymbol{R} \boldsymbol{w} - \mathbf{2} \boldsymbol{w}^T \tilde{\boldsymbol{w}} + 2\lambda \, \|\boldsymbol{w}\|_1. \qquad \text{(Equation S3)}$$

Since $\boldsymbol{R}$ is generally not available in eQTL summary data, $\boldsymbol{R}$ would be approximated by the LD correlation matrix ($\boldsymbol{R}_r$) from an external reference panel such as one from the 1000 Genomes Project[7]. Because $\boldsymbol{R_r}$ and $\tilde{\boldsymbol{w}}$ are derived from genotype data of different samples, the standard LASSO estimates will be unstable and non-unique[3]. The frequentist *lassosum*[3] method replaces $\boldsymbol{R}$ by $\boldsymbol{R}_s = (1-s)\boldsymbol{R_r} + s\boldsymbol{I}$ in Equation S3 with a tuning parameter $0 < s < 1$ and an identity matrix $\boldsymbol{I}$, and then estimates $\hat{\boldsymbol{w}}$ by minimizing the following loss function:

$$f(\boldsymbol{w}) = \frac{\boldsymbol{E}_g^T \boldsymbol{E}_g}{n} + (1-s)\, \boldsymbol{w}^T \boldsymbol{R_r} \boldsymbol{w} - 2\boldsymbol{w}^T \tilde{\boldsymbol{w}} + s\boldsymbol{w}^T \boldsymbol{w} + 2\lambda \|\boldsymbol{w}\|_1. \ \text{(Equation S4)}$$

In our simulation and application studies, pseudovalidation was used to tune $\lambda$ from a grid of 20 equally spaced values from 0.001 to 0.1 on a log-scale and $s$ from (0.2, 0.5, 0.9, 1).

Let $\boldsymbol{X}_r$ denote reference genotype data that are different from the genotype data used to estimate $\tilde{\boldsymbol{w}}$, and $\boldsymbol{R}_r = \boldsymbol{X}_r^T \boldsymbol{X}_r / n_r$ denote the reference LD correlation matrix with reference sample size $n_r$. Given $s$ and $\lambda$, the eQTL effect sizes $\hat{\boldsymbol{w}}$ minimizing $f(\boldsymbol{w})$ in Equation S4 can be obtained by iteratively updating $\hat{w}_j$ as

$$\hat{w}_j^{(k)} = \begin{cases} \dfrac{\operatorname{sign}\left(\mu_j^{(k)}\right)\left|\mu_j^{(k)} - \lambda\right|}{(\tilde{\boldsymbol{X}}_j^T \tilde{\boldsymbol{X}}_j + s)} & \left|\mu_j^{(k)}\right| - \lambda > 0 \\ 0 & \text{otherwise} \end{cases}$$

$$\mu_j^{(k)} = \tilde{w}_j - \tilde{\boldsymbol{X}}_j^T \left( \tilde{\boldsymbol{X}} \hat{\boldsymbol{w}}^{(k-1)} - \tilde{\boldsymbol{X}}_j \, \hat{w}_j^{(k-1)} \right)$$

where $k$ and $k-1$ are the iteration numbers; $\widetilde{\boldsymbol{X}} = \sqrt{1-s}\boldsymbol{X}_r$; and $\widetilde{\boldsymbol{X}}_j$ is the genotype data vector for genetic variant $j$ in $\widetilde{\boldsymbol{X}}$.

To select suitable values for tuning parameters $s$ and $\lambda$, the *lassosum* method uses a *pseudovalidation* procedure that can approximate the correlation between the imputed $\widehat{\boldsymbol{GReX}}$ and the gene expression in the test data using a shrunken estimate of $\widetilde{\boldsymbol{w}}$. We used the *lassosum* package (https://github.com/tshmak/lassosum) to implement the iterative optimization and the *pseudovalidation* procedure. The corresponding proof of iterative optimization and the detailed information of the *pseudovalidation* procedure can be found in the Supplementary materials and the Material and methods section of the *lassosum* paper[3].

#### 3.1.2 Nonparametric Bayesian DPR Method *SDPR*

Following the idea of nonparametric Bayesian DPR model proposed in previous studies[4], the prior of $w_j$ is assumed to be a normal distribution with mean 0 and variance $\sigma_w^2$, with the prior of $\sigma_w^2$ assumed to be a Dirichlet process prior[8]:

$$w_j \sim N(0, \sigma_w^2),\ \sigma_w^2 \sim DP(H, \alpha);$$

where the distribution on $\sigma_w^2$ deviates from the $DP$ with the base distribution $H$ and the concentration parameter $\alpha$ controls the shrinkage of the distribution on $\sigma_w^2$ towards $H$. An equivalent DP normal mixture model can be derived by viewing $\sigma_w^2$ as a latent variable that can be integrated out:

$$w_j | \pi_j, \sigma_j^2 \sim \sum_{j=0}^{+\infty} \pi_j\, N\left(0, \sigma_j^2\right), \quad \pi_j = v_j \prod_{l=0}^{j-1} (1 - v_l), \quad \sigma_j^2 \sim H,$$

$$v_l \sim Beta(1, \alpha),\ \ l = (0, \cdots, j-1, j);$$

The resulting Gaussian mixture prior of $w_j$ is a weighted sum of an infinite number of normal distributions $N(0, \sigma_j^2)$ with weights $\pi_j$ determined by $v_l$ with a Beta prior. The concentration parameter $\alpha$ in the Beta prior distribution determines the number of components with non-zero weights in the mixture normal prior. This infinite Gaussian mixture prior can approximate a wide range of continuous parametric prior distributions assumed by other PRS methods[5,9], for example, the normal distribution in *LDpred*[10], the point normal mixture distribution in *LDpred*[10] and *LDpred2*[11], and the three-point normal mixture distribution in *SBayesR*[12].

*SDPR*[5] implemented DPR but can be applied to summary statistics (i.e., marginal eQTL effect sizes $\widetilde{\boldsymbol{w}}$) as the input by assuming $\widetilde{\boldsymbol{w}}|\boldsymbol{w} \sim N\left(\boldsymbol{Rw}, \frac{\boldsymbol{R}+na\boldsymbol{I}}{n}\right)$, where $a$ is a constant to shrink the covariance between two eQTLs, $\boldsymbol{R}$ is the LD correlation matrix of test eQTLs, and $n$ is the training sample size. The *SDPR* method overparameterizes the model by writing $w_j = \eta\gamma_j$ and use the square of uniform distribution as the base distribution $H$, following Gelman's advice[13]. The *SDPR* method implements a parallel Markov Chain Monte Carlo (MCMC) algorithm in C++ to obtain posterior estimates of $\eta$ and $\gamma_j$ (i.e., $\text{eQTL effect sizes } \widehat{\boldsymbol{w}} = \hat{\eta}\widehat{\boldsymbol{\gamma}}$), conditioning on marginal eQTL effect sizes $\widetilde{\boldsymbol{w}}$ and reference LD correlation matrix $\boldsymbol{R}$. The *SDPR* software also designs an algorithm for partitioning approximately independent LD blocks to ensure that all SNPs in one LD block have no nonignorable correlation with SNPs in other blocks. More details about the MCMC Algorithm and the partition of the reference LD matrix can be found in the Supplemental Text of the *SDPR* paper[5]. For all the parameters, we used the default setting in the *SDPR* software (https://github.com/eldronzhou/SDPR) for our simulation and real studies.

#### 3.1.3 Bayesian Multivariate Regression Method *PRS-CS*

Following the continuous shrinkage prior proposed by *PRS-CS*[6], we assume the following normal prior for $w_j$ and non-informative scale-invariant Jeffreys prior on the residual variance ${\sigma_\epsilon}^2$ in the Equation S1:

$$w_j \sim N\left(0, \frac{\sigma_\epsilon^2}{n}\psi_j\right),\ \ p(\sigma_\epsilon^2) \propto {\sigma_\epsilon}^2;\ \ \psi_j \sim Gamma\big(a, \delta_j\big), \delta_j \sim Gamma(b, \phi),$$

where local shrinkage parameter $\psi_j$ has an independent gamma-gamma prior and $\phi$ is a global-shrinkage parameter controlling the overall sparsity of $\boldsymbol{w}$. Posterior estimates of the eQTL effect sizes $\widehat{\boldsymbol{w}}$ will be obtained by using only summary-level eQTL data $\widetilde{\boldsymbol{w}}$ and the reference LD correlation matrix $\boldsymbol{R}$ by Gibbs Sampler:

$$\boldsymbol{w} \mid \sigma_\epsilon^2, \boldsymbol{\Psi}, \widetilde{\boldsymbol{w}}, \boldsymbol{R} \sim MVN\left(\frac{n}{\sigma_\epsilon^2}\boldsymbol{\Sigma}\widetilde{\boldsymbol{w}}, \boldsymbol{\Sigma}\right), \boldsymbol{\Sigma} = \frac{\sigma_\epsilon^2}{n}(\boldsymbol{R} + \boldsymbol{\Psi}^{-1})^{-1}$$

where $\boldsymbol{\Psi} = \mathrm{diag}\{\psi_1, \psi_2, \cdots, \psi_m\}$. Detailed Gibbs updates of $\sigma_\epsilon^2$, $\psi_j$, and $\delta_j$ in each MCMC iteration has been provided in the Supplemental Note of the *PRS-CS* paper[6].

*PRS-CS* provides two algorithms with different strategies to learn the global shrinkage parameter $\phi$. One is to place a half-Cauchy prior on $\phi$ with $\phi^{1/2} \sim C^{+}(0,1)$, and perform a fully Bayesian approach to learn $\phi$ from the data, which is referred as *PRS-CS auto*. Another is to search a small number of fixed $\phi$ and select the value producing the best predictive performance in the validation data set. In the work, we applied *PRS-CS* with fixed $\phi$, because *PRS-CS auto* tended to fail when the proportion of causal variants is small and the performance of *PRS-CS* is not sensitive to the choice of the fixed $\phi$[6]. Following the suggestion from the *PRS-CS* paper, we set $\phi$ as the square of the proportion of causal variants in the simulation studies and as $10^{-4}$ for each gene in the real data applications. For other parameters, we used the default setting in the *PRS-CS* software (https://github.com/getian107/PRScs).

#### 3.2 OTTERS Stage II Gene-based Association Test

Once we train eQTL weights by using each training method from the summary-level reference eQTL data, we will perform the respective gene-based association analysis for that method by taking the trained eQTL weights as corresponding SNP weights. We thus derive a set of TWAS p-values per gene g; one p-value for each training method that we applied. Finally, we will employ the aggregated Cauchy association test (ACAT-O) approach[14] to combine the TWAS p-values from individual training methods and conduct an omnibus test. We refer to these p-values obtained by ACAT-O tests as OTTERS p-values. We describe how we perform the gene-based association analysis and construct the OTTERS p-values below.

##### 3.2.1 Gene-based Association Test with eQTL Weights

Given individual-level GWAS data with genotype $\boldsymbol{X}_{new}$ and phenotype $\boldsymbol{Y}$, the GReX for $\boldsymbol{X}_{new}$ can be imputed by $\widehat{\boldsymbol{GReX}} = \boldsymbol{X}_{new}\, \widehat{\boldsymbol{w}}$. The follow-up TWAS using a gene-based burden testing is to assess the association between $\widehat{\boldsymbol{GReX}}$ and $\boldsymbol{Y}$ based on the following generalized linear regression model:

$$f(E[\boldsymbol{Y}|\boldsymbol{X_{new}}, \boldsymbol{C}] = \theta \boldsymbol{C} + \beta\, \widehat{\boldsymbol{GReX}},$$

where $\boldsymbol{C}$ denotes the matrix for covariates associated with $\boldsymbol{Y}$, and $f(\cdot)$ denotes a pre-specified link function. This link function can be set as identity function for quantitative phenotype and logit function for dichotomous phenotype. The gene-based association test is equivalent to test $H_0: \beta = 0$.

When summary-level GWAS data are available, one can use the TWAS Z-score test statistics as used by FUSION[15] and S-PrediXcan[16] as follows:

$$Z_{g,FUSION} = \frac{\sum_{j=1}^{J}(\hat{w}_j Z_j)}{\sqrt{\hat{\boldsymbol{w}}'\boldsymbol{V}\hat{\boldsymbol{w}}}}, \quad Z_{g,S-PrediXcan} = \frac{\sum_{j=1}^{J}(\hat{w}_j \hat{\sigma}_j Z_j)}{\sqrt{\hat{\boldsymbol{w}}'\boldsymbol{V}\hat{\boldsymbol{w}}}};$$

where $Z_j$ is the single variant Z-score test statistic in GWAS for the j[th] SNP, $j = 1, \dots, \mathrm{J}$, for all test SNPs that have both eQTL weights with respect to the test gene $g$ and GWAS Z-scores; $\hat{\sigma}_j$ is the genotype standard deviation of the j[th] SNP; and $\boldsymbol{V}$ denotes the genotype correlation matrix in FUSION Z-score statistic and genotype covariance matrix in S-PrediXcan Z-score statistic of the test SNPs.

The Supplementary Text of our previous work TIGAR-V2[17] has proved that when $\hat{\boldsymbol{w}}$ are estimated using standardized reference data, FUSION Z-score statistic is equivalent to the SPrediXcan test statistic; while when $\hat{\boldsymbol{w}}$ are estimated using non-standardized reference data, FUSION Z-score statistic will lead to inflated Type I error and SPrediXcan test statistic should be used in this situation. In this work, we obtained standardized $\hat{\boldsymbol{w}}$ from the *lassosum*, *SDPR*, and *PRS-CS* software. For *P+T*, we estimated the standardized effect size based on the Z-score statistic values in the eQTL summary data by $\tilde{w}_j \approx Z_j / \sqrt{median(n_{g,j})}$, where $median(n_{g,j})$ denotes the median sample size of all cis-eQTLs for the target gene $g$.

#### 3.2.2 OTTERS p-value by ACAT-O Test

Following the idea of ACAT-O[14] to combine the TWAS p-values using eQTL weights trained by different methods from the summary-level reference eQTL data, we create our OTTERS test statistics as:

$$T_{OTTERS} = \frac{1}{K}\sum_{k=1}^{K} \tan\{(0.5 - p_k)\}\pi\} \ , \qquad \pi \approx 3.14;$$

where $p_k$ is the TWAS p-value with eQTL weights trained using the $k$th method, e.g., *P+T(0.001), P+T(0.05), lassosum, SDPR,* and *PRS-CS* (K=5). Under null hypothesis, the p-value $p_k$ is uniformly distributed in $[0, 1]$ and the transformation $\tan\{(0.5 - p_k)\}$ is Cauchy distributed. Thus, the null distribution of the test statistic $T_{OTTERS}$ can be approximated by a Cauchy distribution[18] and the corresponding OTTERS p-value ($p_{OTTERS}$) can be approximated by

$$p_{OTTERS} \approx \frac{1}{2} - \frac{\{\arctan(T_{OTTERS})\}}{\pi}.$$
